## Supplementary Material for "SpikeMAP: An unsupervised pipeline for the identification of cortical excitatory and inhibitory neurons in high-density multielectrode arrays with ground-truth validation"

**Supplemental Figures**

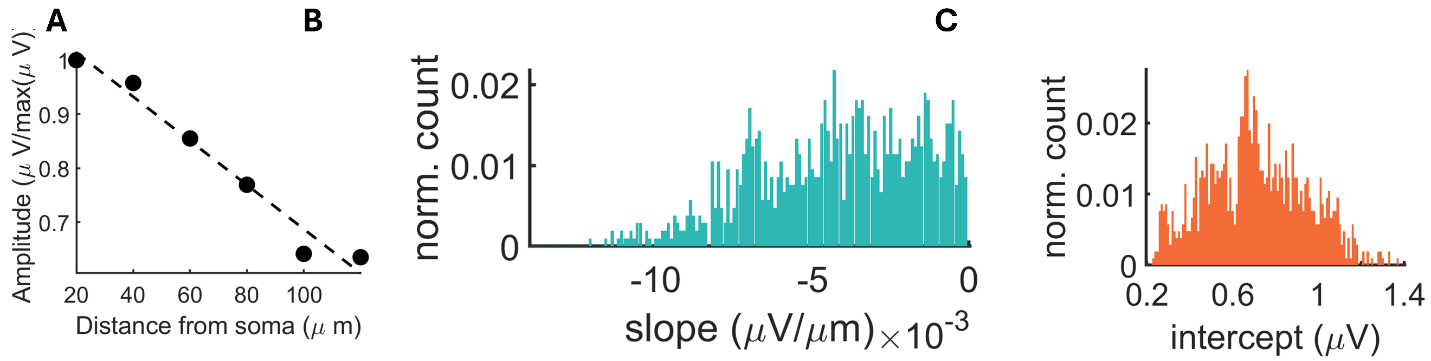

****Supplemental Figure 1:** (**A**) Voltage amplitude as a function of distance from highest electrode peak. Putative somas with an increase in voltage away from electrode peak were excluded from this analysis. (**B-C**) Distribution of slopes and intercepts across all putative somas obtained from one recording (*N*=1,950).**

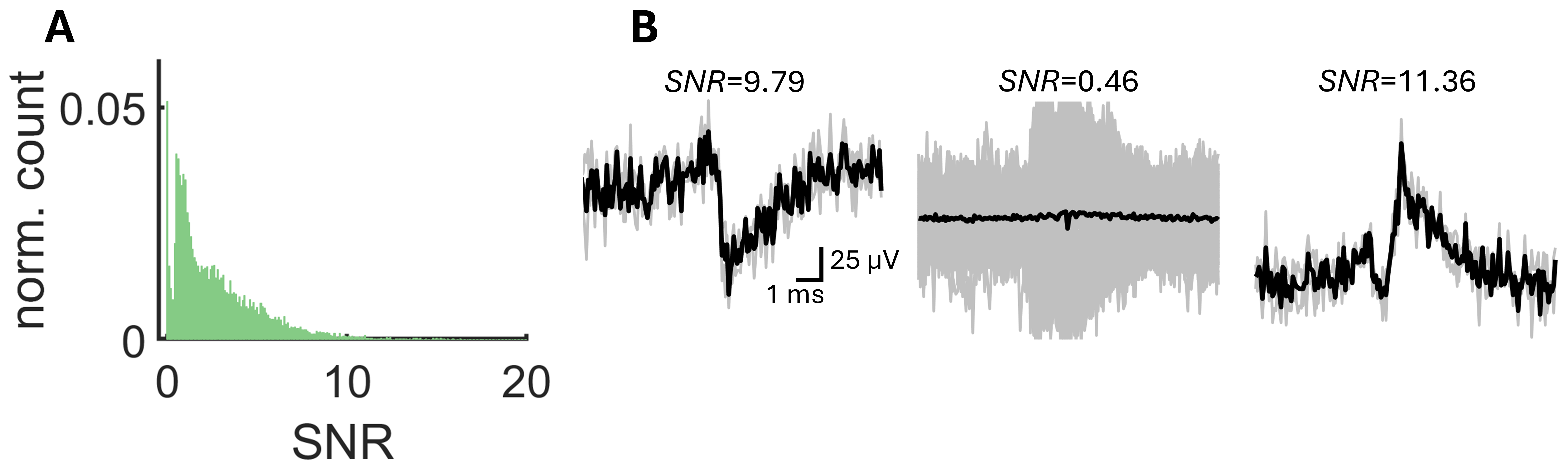

****Supplemental Figure 2:** (**A**) Distribution of signal-to-noise ratio (SNR) over all spike-sorted clusters of a single recording (*N*=8,059). (**B**) Examples of mean voltages (black lines) and individual voltages (grey lines) around individual spikes.**

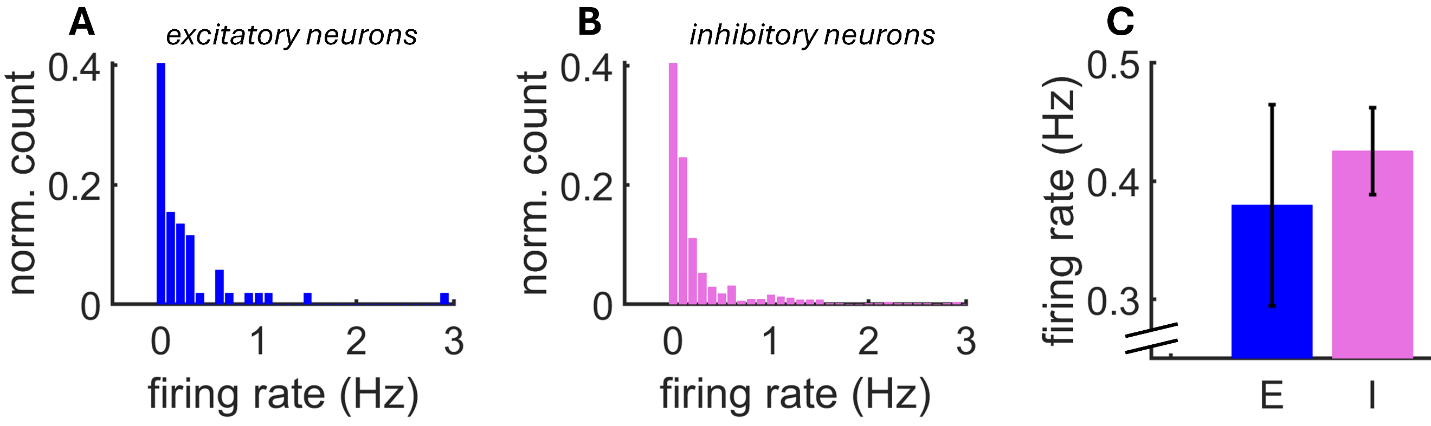

****Supplemental Figure 3:** Distribution of mean firing rates across a population of putative excitatory (**A**) (*N*=570) and inhibitory (**B**) (*N*=54) neurons. (**C**) Mean firing rate across neurons. Vertical bars: SEM.**

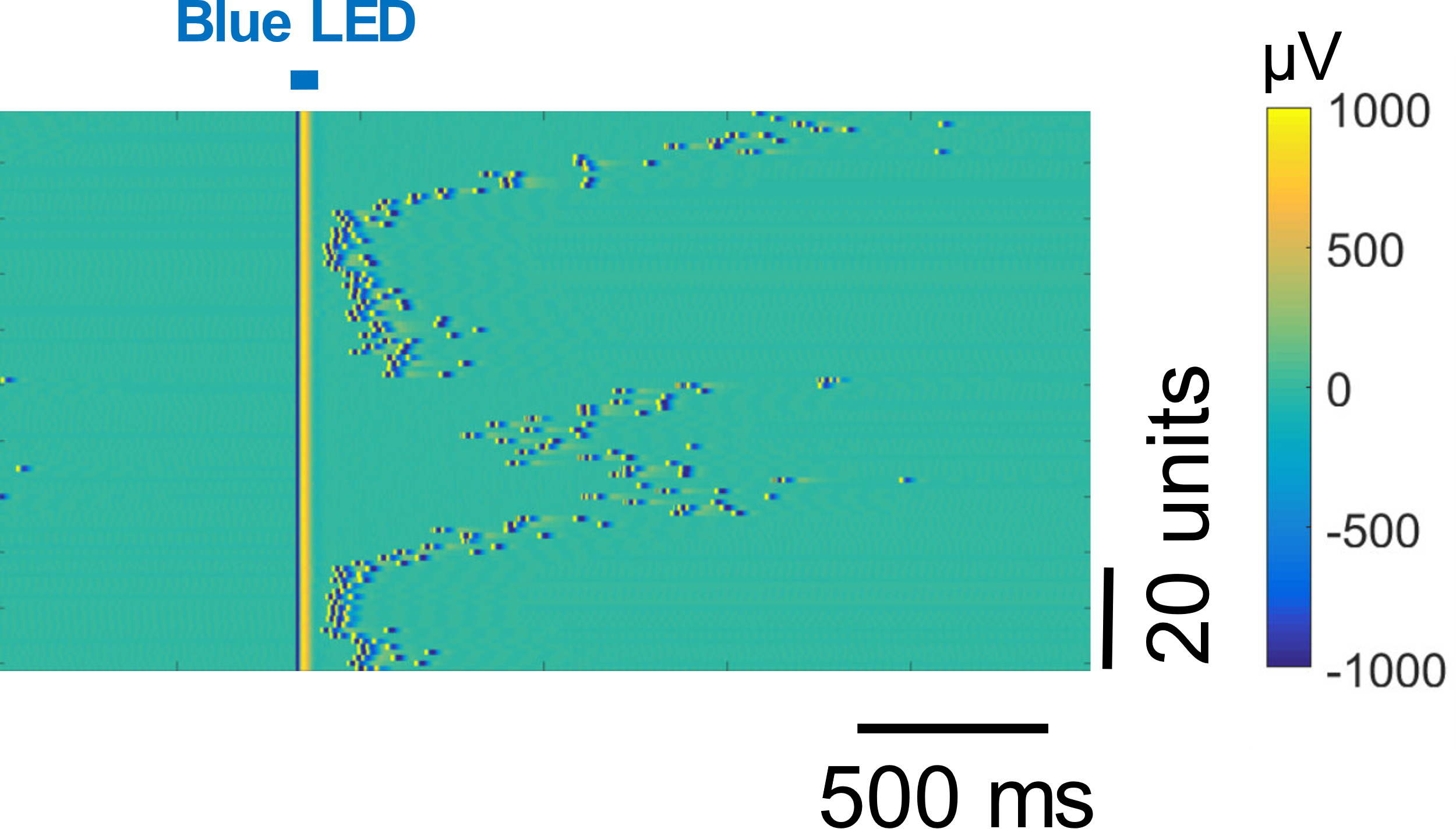

**Supplemental Figure 4:** Light artefact in a recording from mouse prefrontal cortex showing that the application of a brief (1 ms) pulse of light leads to saturation lasting about 5 ms post-stimulation.

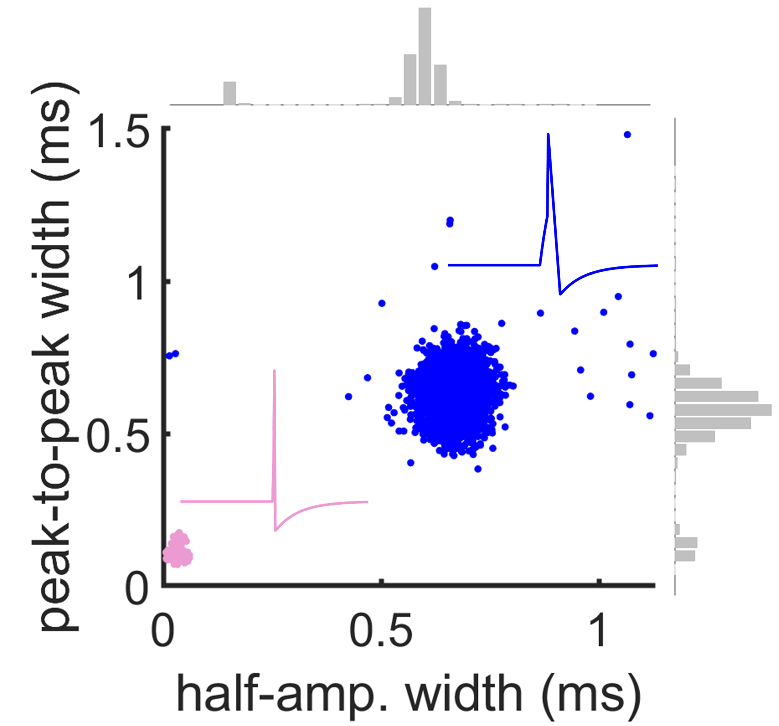

****Supplemental Figure 5:**** Waveform kinetics for E and I cells obtained during the baseline window of the optogenetic protocol. Insets show cartoons of individual spikes with broader (E) and narrower (I) waveforms.

| File name | Animal # | Protocol # | # of E cells  identified | # of I cells  identified | total recording time (min) |
| --- | --- | --- | --- | --- | --- |
| R20211221_Slice1_Vars3.mat | 1 | 1 | 3687 | 47 | 8 |
| R20211213_Slice2_Vars3.mat | 2 | 1 | 2943 | 350 | 8 |
| R20211209_Slice3_Vars3.mat | 3 | 1 | 1994 | 712 | 8 |
| R20211219_Slice3_Vars3.mat | 4 | 2 | 1138 | 32 | 8 |

****Supplemental Table 1:** Summary of SpikeMAP open-source data. Datasets were obtained from four mice. The optogenetic stimulation protocol (see Materials and Methods) was run once per dataset. Data is available from: dx.doi.org/10.6084/m9.figshare.29416472**
